# The Fission Yeast S-Phase Cyclin Cig2 Can Drive Mitosis

**DOI:** 10.1101/213330

**Authors:** Mira Magner, Mary Pickering, Dan Keifenheim, Nicholas Rhind

**Author notes:** Present address: Department of Genetics, Cell Biology and Development, University of Minnesota, 420 Washington Avenue SE, Minneapolis MN 55455 USA.

## Abstract

Commitment to mitosis is regulated by cyclin-dependent kinase (CDK) activity. In the fission yeast *Schizosaccharomyces pombe*, the major B-type cyclin, Cdc13, is necessary and sufficient to drive mitotic entry. Furthermore, Cdc13 is also sufficient to drive S phase, demonstrating that a single cyclin can regulate alternating rounds of replication and mitosis and providing the foundation of the quantitative model of CDK function. It has been assumed that Cig2, a B-type cyclin expressed only during S-phase and incapable of driving mitosis in wild-type cells, was specialized for S-phase regulation. Here, we show that Cig2 is capable of driving mitosis. Cig2/CDK activity drives mitotic catastrophe—lethal mitosis in inviably small cells—in cells that lack CDK inhibition by tyrosine-phosphorylation. Moreover, Cig2/CDK can drive mitosis in the absence of Cdc13/CDK activity and constitutive expression of Cig2 can rescue loss of Cdc13 activity. These results demonstrate that in fission yeast, not only can the presumptive M-phase cyclin drive S phase, but the presumptive S-phase cyclin can drive M phase, further supporting the quantitative model of CDK function. Furthermore, these results provide an explanation, previously proposed on the basis of computational analyses, for the surprising observation that cells expressing a single-chain Cdc13-Cdc2 CDK do not require Y15 phosphorylation for viability. Their viability is due to the fact that in such cells, which lack Cig2/CDK complexes, Cdc13/CDK activity is unable to drive mitotic catastrophe.

## Introduction

Cdc13, the major B-type cyclin in the fission yeast *Schizosaccharomyces pombe*, binds to and activates the Cdc2 cyclin-dependent kinase (CDK) (Booher et al., 1989; Moreno et al., 1989). Of the five characterized fission yeast cyclins, Cdc13 alone is necessary to drive mitotic entry and sufficient to drive essentially normal cell cycles (Fisher and Nurse, 1995; Fisher and Nurse, 1996). Cig2 is another B-type cyclin (Bueno and Russell, 1993). It is expressed during S-phase and was thought to be limited to S-phase functions because, although Cig2 is capable of driving S-phase, it is unable to drive mitosis in wild-type cells (Martin-Castellanos et al., 1996; Mondesert et al., 1996).

The fact that a single B-type cyclin is sufficient for normal cell-cycle control in fission yeast has been instrumental in the development of the quantitative model of CDK function. This model posits that the difference between the CDK activities that trigger S phase and M phase is quantitative—that the same CDK activity can trigger both, but that a lower level of activity is required for entry into S phase than is required for entry into M (Fisher and Nurse, 1996; Stern and Nurse, 1996). Such a model explains why fission yeast needs only one cyclin and, more generally, explains why S phase precedes M phase in the cell cycle. In its simplest form, the model envisions three levels of B-type cyclin/CDK (B/CDK) activity: OFF, the level during G1 when B-type cyclins are not expressed and B-type-specific CDK inhibitors (CKIs) are expressed; LOW, the level of activity that is sufficient to trigger S phase, but not mitosis, and which is provided by B/CDK inhibited by tyrosine-15 (Y15) phosphorylation; and HIGH, the level provided by Y15-dephosphorylated B/CDK, which is required to trigger mitosis. It should be noted that the basic quantitative model deals only with the B/CDK activity that regulate the S and M phases of the cell cycle. It does not address the G1/CDK activity that is required for exiting G1 and initiating the cell cycle (Futcher, 1996).

An alternate explanation for the ordering of cell-cycle events is the qualitative model of CDK function, which posits that qualitatively different cyclin/CDK complexes phosphorylate different substrates to trigger S and M phase (Cross et al., 1999; Loog and Morgan, 2005). Although modern eukaryotic cell cycles have aspects of both the quantitative and qualitative models (Uhlmann et al., 2011), a plausible argument can be made that the original eukaryotic cell cycle was closer to the quantitative model and that qualitative aspects have been added to cell cycle regulation as eukaryotes have diversified (Swaffer et al., 2016).

Support for the quantitative model was provided by an elegant study in which Cdc13 and Cdc2 were expressed as a translationally-fused, single-chain CDK, removing the possibility of other Cdc2 cyclin/CDK complexes forming. The single CDK is able to drive normal mitotic and meiotic cell cycles (Coudreuse and Nurse, 2010; Gutiérrez-Escribano and Nurse, 2015), demonstrating that qualitatively different CDK activity is not required for ordered cell-cycle progression. An unexpected result from this work was that the single-chain-expressing cells do not require Y15-phosphorylation-dependent CDK regulation for viability or proper size control. Inhibitory Y15 phosphorylation of CDKs is a critical point of eukaryotic cell cycle regulation (Nurse, 1990). In the fission yeast, as in most other eukaryotes that have been studied, dephosphorylation of Cdc2 Y15 is the rate-limiting step for entry into mitosis (Rhind and Russell, 2012) and thus critical for preventing premature mitosis and regulating the size of cells at mitosis. Loss of the Wee1 and Mik1 kinases, which catalyze Y15 phosphorylation, leads to mitotic catastrophe, in which cells enter mitosis at an inviably small size (Lundgren et al., 1991). Since Cdc13 is necessary and sufficient for mitotic entry during normal fission yeast cell cycles, it has been presumed to be the cyclin responsible for driving cells into mitosis during the mitotic catastrophe caused by loss of Y15 phosphorylation. The fact that the Cdc13-Cdc2 single-chain CDK does not require Y15 phosphorylation led to speculation that there may be other, Y15-independent, mechanisms of CDK regulation at the G2/M transition (Coudreuse and Nurse, 2010; Navarro and Nurse, 2012). However, as pointed out by a recent computational analysis (Gerard et al., 2015), if other cyclins, such as Cig2, contribute to the mitotic catastrophe phenotype, the viability of the single-chain CDK cells in the absence of Y15 phosphorylation can be explained without the need for additional mechanisms of CDK regulation. Instead their viability could be explained by the fact that they lack Cig2/CDK activity, which, in this scenario, would be responsible for the inviability of cells lacking Y15 phosphorylation.

To explore the regulation of fission yeast CDK, we have tested the ability of Cig2 to drive mitosis. Cig2/CDK is required for mitotic catastrophe in the absence of Cdc2 Y15 phosphorylation (Bueno and Russell, 1993; Zarzov et al., 2002). We find that Cig2/CDK activity is not only necessary for mitotic catastrophe, but is also sufficient to drive mitosis in the absence of Cdc13/CDK activity. Our results show that Cig2, although involved primarily in S-phase regulation in wild-type cells, is capable of driving mitosis, consistent with the quantitative model of CDK activity. Furthermore, the requirement of Cig2 for mitotic catastrophe in the absence of Cdc2 Y15 phosphorylation explains the viability of Cdc13-Cdc2 single-chain CDK cells lacking Y15 phosphorylation without the need for further mechanisms of CDK regulation.

## Results

We observed that Cig2 is required for the lethal mitosis that occurs when Cdc2 is prematurely activated by removal of the Wee1 and Mik1 tyrosine phosphatases (Figure 1). *wee1-50ts* is a strong temperature sensitive allele of *wee1* that retains less than 10% activity *in vivo* at 30°C (Rhind and Russell, 2001). *wee1-50ts mik1Δ* cells rapidly dephosphorylate Cdc2 and enter mitosis when shifted to 30°C, regardless of cell size (Lundgren et al., 1991 and Figure 1). Multiple rounds of rapid division without sufficient growth leads to inviably small cells, a phenotype referred to as mitotic catastrophe (Lundgren et al., 1991). This synthetic lethality between *wee1-50ts* and *mik1Δ* at 30°C is suppressed by loss of Cig2 (Bueno and Russell, 1993 and Figure 1). *wee1-50ts mik1Δ cig2Δ* cells are viable at 30°C, but divide at the short length of 8.4±1.3 *μ*m (Figures 1B, 1C and Table 1). We interpret this suppression to indicate that the lethal mitoses, which occur in the smallest cells, require Cig2/CDK activity instead of, or in addition to, Cdc13/CDK activity.

**Table 1.**
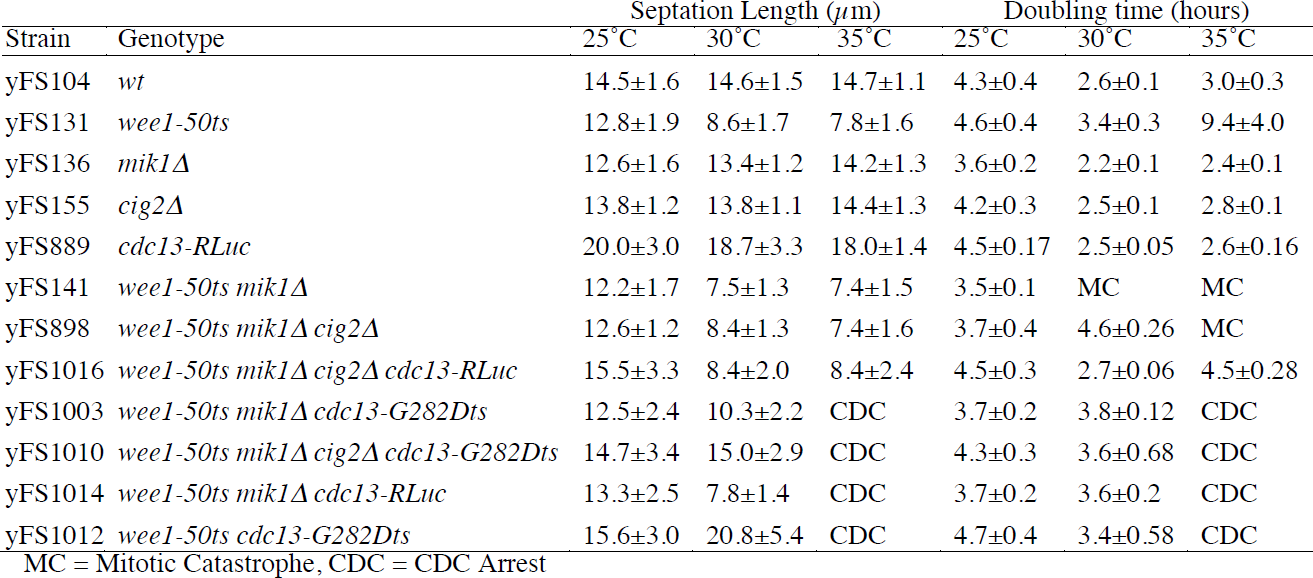
Cell lengths at division and doubling times

| Strain | Genotype | Septation Length ($\mu\text{m}$) | | | Doubling time (hours) | | |
| --- | --- | --- | --- | --- | --- | --- | --- |
|  |  | 25°C | 30°C | 35°C | 25°C | 30°C | 35°C |
| yFS104 | *wt* | 14.5±1.6 | 14.6±1.5 | 14.7±1.1 | 4.3±0.4 | 2.6±0.1 | 3.0±0.3 |
| yFS131 | *wee1-50ts* | 12.8±1.9 | 8.6±1.7 | 7.8±1.6 | 4.6±0.4 | 3.4±0.3 | 9.4±4.0 |
| yFS136 | *mik1Δ* | 12.6±1.6 | 13.4±1.2 | 14.2±1.3 | 3.6±0.2 | 2.2±0.1 | 2.4±0.1 |
| yFS155 | *cig2Δ* | 13.8±1.2 | 13.8±1.1 | 14.4±1.3 | 4.2±0.3 | 2.5±0.1 | 2.8±0.1 |
| yFS889 | *cdc13-RLuc* | 20.0±3.0 | 18.7±3.3 | 18.0±1.4 | 4.5±0.17 | 2.5±0.05 | 2.6±0.16 |
| yFS141 | *wee1-50ts mik1Δ* | 12.2±1.7 | 7.5±1.3 | 7.4±1.5 | 3.5±0.1 | MC | MC |
| yFS898 | *wee1-50ts mik1Δ cig2Δ* | 12.6±1.2 | 8.4±1.3 | 7.4±1.6 | 3.7±0.4 | 4.6±0.26 | MC |
| yFS1016 | *wee1-50ts mik1Δ cig2Δ cdc13-RLuc* | 15.5±3.3 | 8.4±2.0 | 8.4±2.4 | 4.5±0.3 | 2.7±0.06 | 4.5±0.28 |
| yFS1003 | *wee1-50ts mik1Δ cdc13-G282Dts* | 12.5±2.4 | 10.3±2.2 | CDC | 3.7±0.2 | 3.8±0.12 | CDC |
| yFS1010 | *wee1-50ts mik1Δ cig2Δ cdc13-G282Dts* | 14.7±3.4 | 15.0±2.9 | CDC | 4.3±0.3 | 3.6±0.68 | CDC |
| yFS1014 | *wee1-50ts mik1Δ cdc13-RLuc* | 13.3±2.5 | 7.8±1.4 | CDC | 3.7±0.2 | 3.6±0.2 | CDC |
| yFS1012 | *wee1-50ts cdc13-G282Dts* | 15.6±3.0 | 20.8±5.4 | CDC | 4.7±0.4 | 3.4±0.58 | CDC |

MC = Mitotic Catastrophe, CDC = CDC Arrest

**Figure 1.**
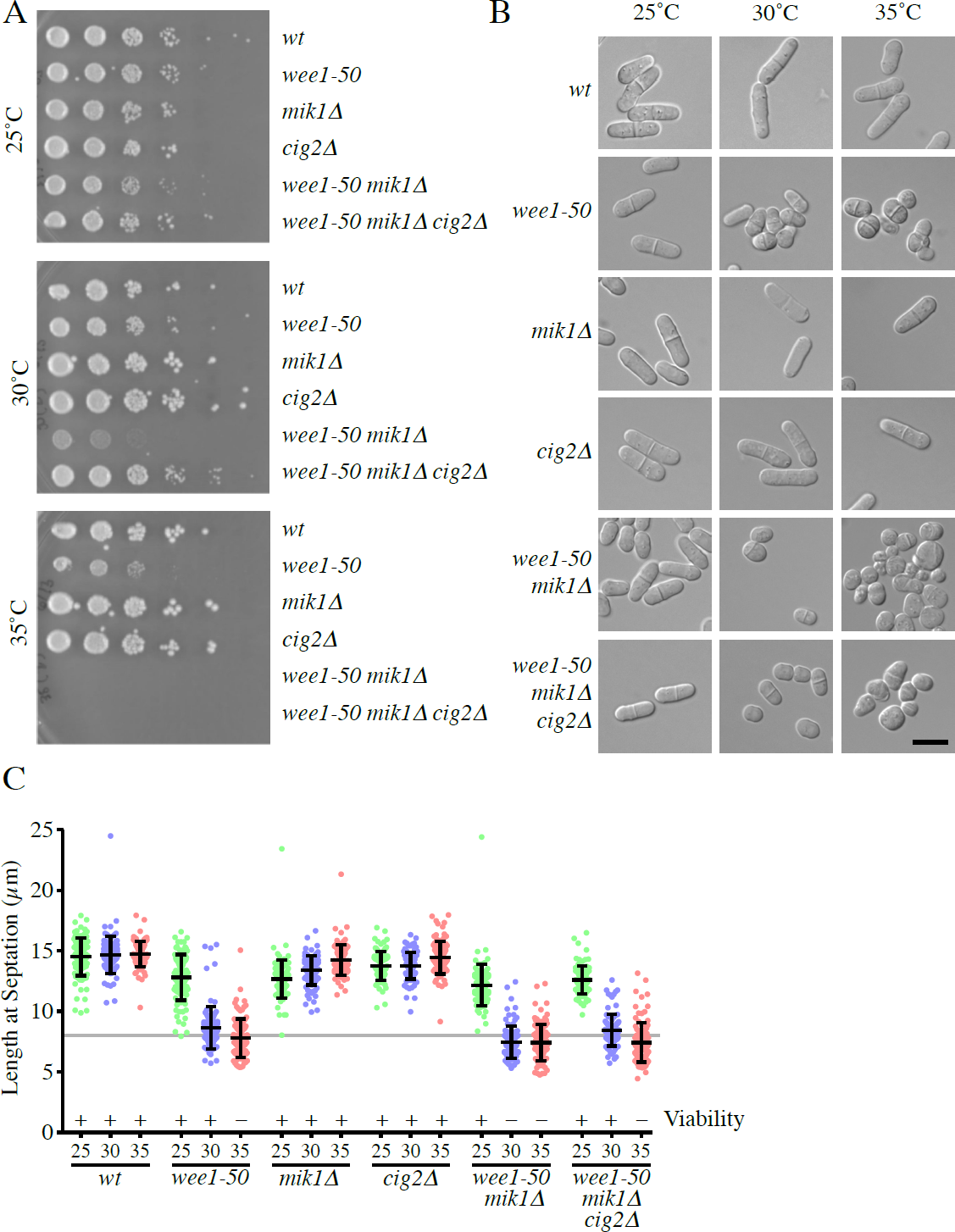
cig2Δ suppresses the mitotic catastrophe phenotype of wee1-50ts mik1Δ cells. A) Dilution spot assay of cell viability. yFS104 (wild type), yFS131 (*wee1-50ts*), yFS136 (*mik1Δ*), yFS155 (*cig2Δ*), yFS141 (*wee1-50ts mik1Δ*) and yFS898 (*wee1-50ts mik1Δ cig2Δ*) cells were grown to mid-log phase at 25°C in YES, serially 10-fold diluted, spotted onto YES plates and grown for three days at the indicated temperature. B) DIC micrographs of the same strains grown to mid-log phase at the indicated temperature in liquid YES. The scale bar is 10 *μ*m. C) Cells from the cultures in B were photographed and measured for length at septation. Error bars represent mean and standard deviation. n = 100.

To test if loss of Cig2 could suppress the complete loss of Cdc2 Y15 phosphorylation, we attempted, but were unable, to build a *wee1Δ mik1Δ cig2Δ* strain. Consistent with the conclusion that loss of *cig2* is unable to suppress complete loss of *wee1* and *mik1*, we found that *wee1-50ts mik1Δ cig2Δ* are inviable at 35°C, a temperature at which *wee1-50ts* retains less than 0.5% activity (Rhind and Russell, 2001). From these results, we infer that the residual Wee1 activity in *weel-50* cells at 30°C is sufficient to inhibit the low levels of Cdc13/CDK present during S-phase, but at 35°C it is not, and the low level of Cdc13/CDK present can drive cells into lethal premature mitosis. We note that strains with an average length at septation above about 8 *μ*m are viable, whereas those that divide below 8 *μ*m on average are not, suggesting that 8 *μ*m is about the minimal size for viable cell division in *S. pombe* and that Cig2 can drive cells into mitosis below that viable limit in situations in which Cdc13 alone cannot.

To confirm that Cig2 can drive mitosis at smaller cell sizes than Cdc13, we examined the kinetics of mitotic entry in synchronized cells. We find that loss of Cig2 delays the entry into mitosis when small *wee1-50ts mik1Δ* cells are shifted to 30°C, such that *wee1-50ts mik1Δ* cells divide at a smaller size than *wee1-50ts mik1Δ cig2Δ* cells in synchronous cultures (8.8±0.8 *μ*m v. 9.9±0.9 *μ*m) (Figure 2A). To be able to determine more accurately how quickly after loss of Cdc2 tyrosine phosphorylation cells of different sizes can divide, we also filmed asynchronous cultures as they were shifted to 30°C. Again, we find that *wee1-50ts mik1Δ* cells can divide at a smaller size than *wee1-50ts mik1Δ cig2Δ* cells (9.8±0.9 *μ*m v. 10.8±0.9 *μ*m) (Figure 2B). The one micron difference in results between the elutriation and time-lapse measurements is likely due to the microscope stage being at a lower temperature than the target of 30°C. We interpret these results to indicate that, in small cells, there is insufficient Cdc13 to activate mitosis, and that mitotic catastrophe in *wee1-50ts mik1Δ* cell is driven by Cig2. However, as cells grow, they accumulate sufficient Cdc13 to activate mitosis in the absence of Cig2.

**Figure 2.**
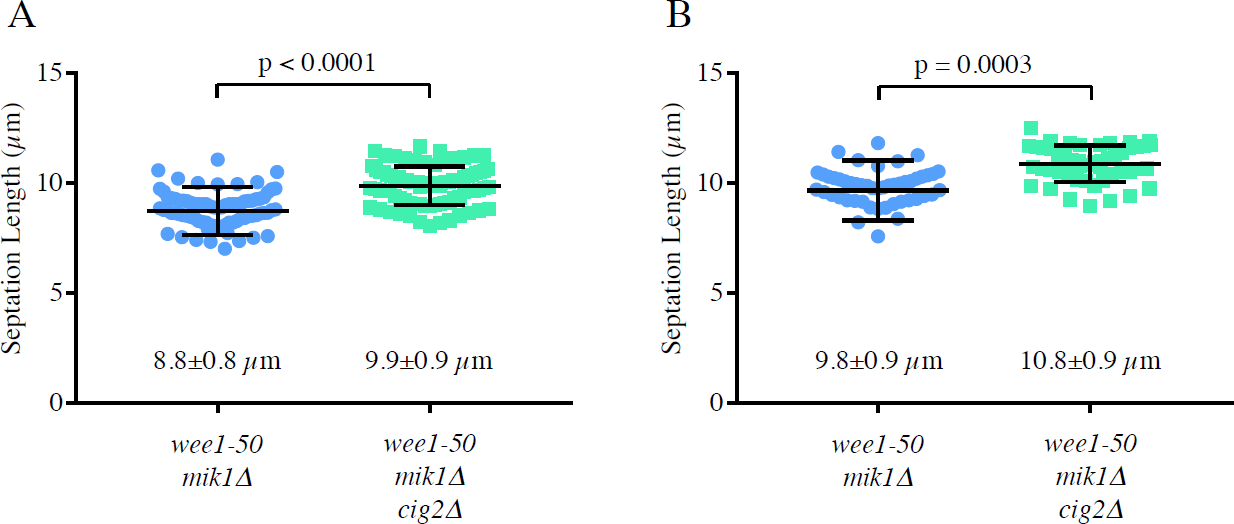
Cig2 drives mitosis at a smaller cell size than Cdc13. A) yFS141 (*wee1-50ts mik1Δ*) and yFS898 (*wee1-50ts mik1Δ cig2Δ*) cells were grown to mid-log phase at 25°C, synchronized in late S phase/early G2 by elutriation and immediately shifted to 30°C. Cells were sampled every 20 minutes, photographed, and measured for length at septation. B) yFS141 (*wee1-50ts mik1Δ*) and yFS898 (*wee1-50ts mik1Δ cig2Δ*) cells were grown to mid-log phase at 25°C, transferred to a microscope stage at 30°C, filmed and measured for length at septation. Statistical significance of the difference in distribution of lengths at septation was calculated with Student’s t test. n = 100.

The conclusion that Cig2 and Cdc13 collaborate to induce mitotic catastrophe in *wee1-50ts mik1Δ* cells at 35°C suggests that the amount of Cdc13 present during S-phase is only just sufficient to drive entry into mitosis in the absence of Cdc2 Y15 phosphorylation. To test this possibility, we took advantage of the fact that a C-terminal *Renilla* luciferase tag on Cdc13 slightly reduces Cdc13/CDK activity, creating a hypomorphic *cdc13* allele (Table 1). We built a *wee1-50ts mik1Δ cig2Δ cdc13-RLuc* strain and found that it is viable at 35°C (Figure 3 and Table 1), suggesting that the lethal mitoses observed in *wee1-50ts mik1Δ cig2Δ* cells at 35°C is due to early accumulation of Cdc13/CDK activity that is reduced in the *cdc13-RLuc* allele. Consistent with this interpretation, *wee1-50ts mik1Δ cdc13-RLuc* cells are viable at 30°C (Figure 3). These results suggest that Cig2 and Cdc13 both contribute to the CDK activity responsible for driving lethal mitoses in *wee1-50ts mik1Δ* cells at 30°C and the removal of either is sufficient to prevent mitotic catastrophe.

**Figure 3.**
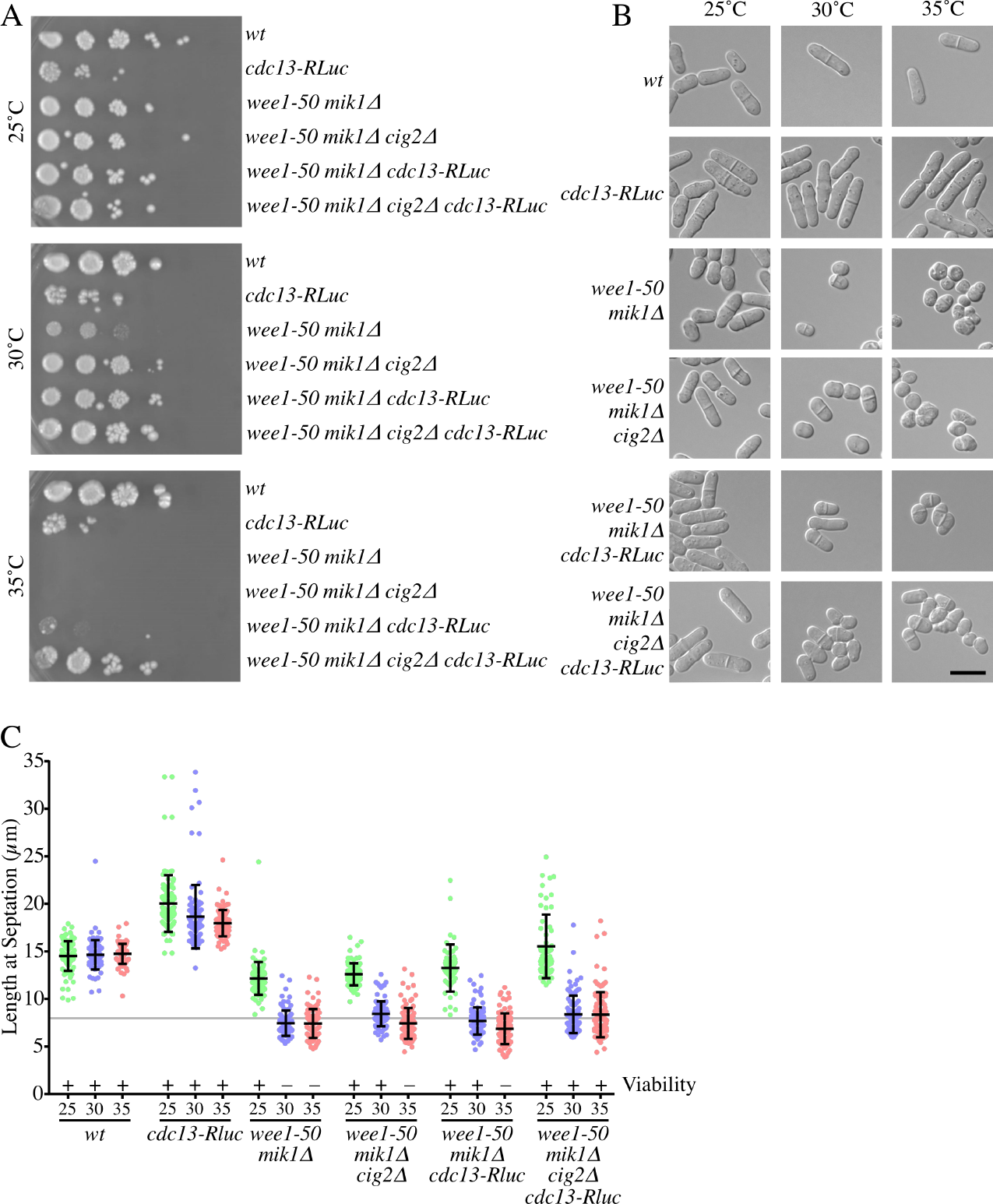
Cig2 and Cdc13 both contribute to premature mitosis in *wee1-50ts mik1Δ* cells. A) Dilution spot assay of cell viability. yFS104 (wild type), yFS889 (*cdc13-RLuc*), yFS141 (*wee1-50ts mik1Δ*), yFS898 (*wee1-50ts mik1Δ cig2Δ*), yFS1014 (*wee1-50ts mik1Δ cdc13-RLuc*) and yFS1016 (*wee1-50ts mik1Δ cig2Δ cdc13-RLuc*) cells were grown to mid-log phase at 25°C in YES, serially 10-fold diluted, spotted onto YES plates and grown for three days at the indicated temperatures. B) DIC micrographs of the same strains grown to mid-log phase at the indicated temperature in liquid YES. The scale bar is 10 *μ*m. C) Cells from the cultures in B were photographed and measured for length at septation. Error bars represent mean and standard deviation. n = 100.

To test if Cig2 is involved in mitosis in normal cell cycles, we examined the genetic interaction between *cig2Δ* and *cdc13-G282Dts*, a strong, temperature-sensitive allele of *cdc13*, which shows a nearly null phenotype at 36°C (Berry and Gould, 1996). We find that loss of Cig2 sensitizes *cdc13-G282Dts* cells, reducing their restrictive temperature and increasing the severity of their terminal phenotype (Figure 4). In particular, the double mutant has both more mitotic abnormalities at the permissive temperature of 25°C and a tighter arrest phenotype at the restrictive temperature of 36°C, with fewer cells leaking into an abnormal mitosis (Figure 4C). These results further demonstrate that Cig2 can contribute to mitotic progression in cells compromised for Cdc13/CDK activity.

**Figure 4.**
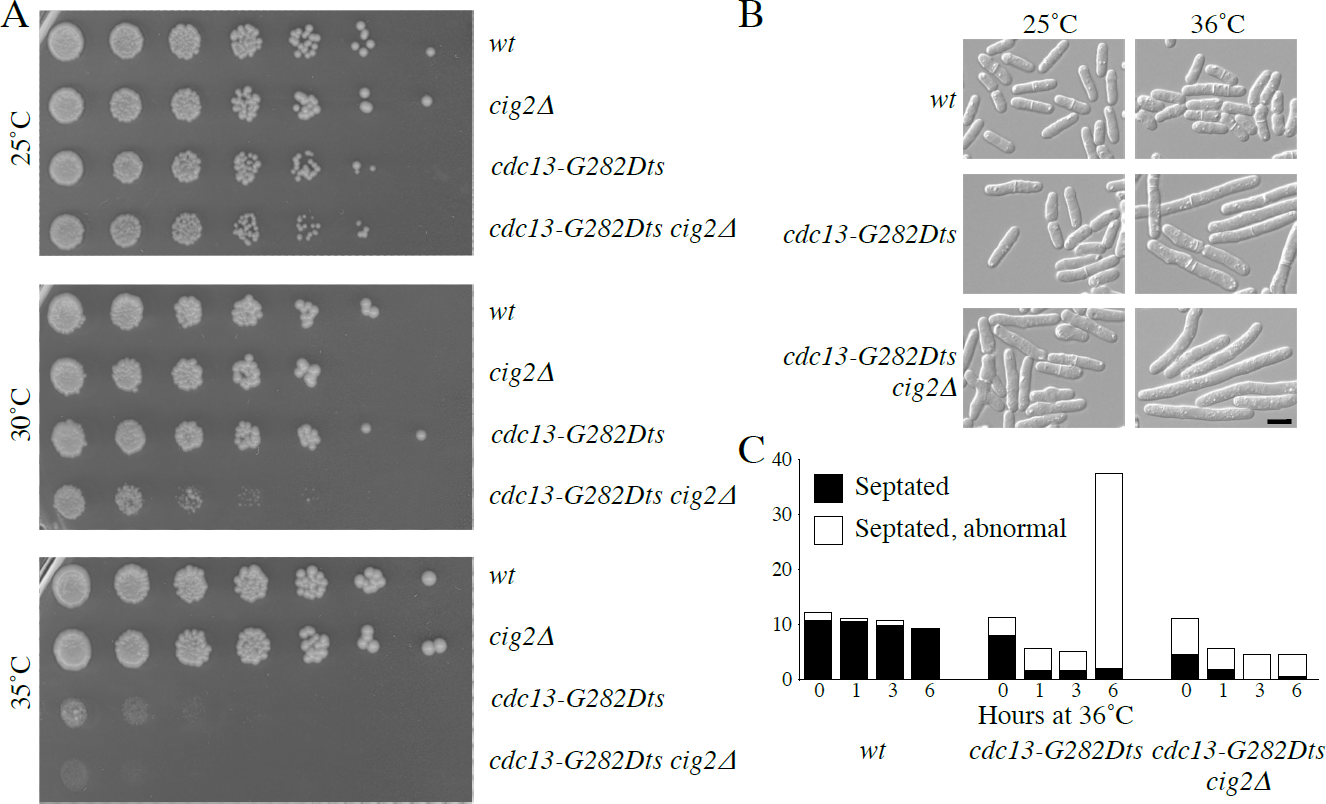
Cig2 promotes mitoses during normal cell cycles. A) Dilution spot assay of cell viability. yFS104 (wild type), yFS155 (*cig2Δ*), yFS1032 (*cdc13-G282Dts*) and yFS1046 (*cdc13-G282Dts cig2Δ*) cells were grown to mid-log phase at 25°C in YES, serially 4-fold diluted, spotted onto YES plates and grown for three days at the indicated temperatures. B) DIC micrographs of the same strains grown to mid-log phase in liquid YES at 25°C and incubated at the indicated temperature for 6 hours. The scale bar is 10 *μ*m. C) Cells from the cultures in B were photographed and counted for septation. Abnormal septation includes multiple septa, misshapen septa and asymmetric septation. n > 200.

Having shown that Cig2 can be involved in driving cells into mitosis, we were interested to know if Cig2 is sufficient to drive entry into mitosis is the absence of Cdc13. To test this possibility, we examined mitosis in *wee1-50ts mik1Δ cdc13-G282Dts* cells, which harbor strong, temperature-sensitive alleles of *wee1* and *cdc13*, both of which show nearly null phenotypes at 35˚C (Berry and Gould, 1996; Rhind and Russell, 2001). We reasoned that by inactivating both Wee1 and Cdc13 at the same time, we would be able to assay the ability of Cig2 to drive mitosis in the absence of Cdc13. *wee1-50ts mik1Δ cdc13-G282Dts* cells shifted to 35°C immediately after elutriation synchronization—a time when most cells are in late S phase or early G2—proceed rapidly into mitosis. This transition is dependent upon Cig2, as demonstrated by the fact that >95% of *wee1-50ts mik1Δ cdc13-G282Dts cig2Δ* cells shifted at the same time arrest in G2 (Figure 5B). Most *wee1-50ts mik1Δ cdc13-G282Dts* cells that divide at 35°C display a morphologically normal mitosis and cytokinesis, although a minority show defects, such as multiple or asymmetric septa (Figure 5A). Moreover, most *wee1-50ts mik1Δ cdc13-G282Dts* cells that divide at 35°C are viable when shifted back to 25°C after division (Figure 5C). These results show that Cig2 is able to drive a functional mitosis in cells lacking most or all Cdc13 activity.

**Figure 5.**
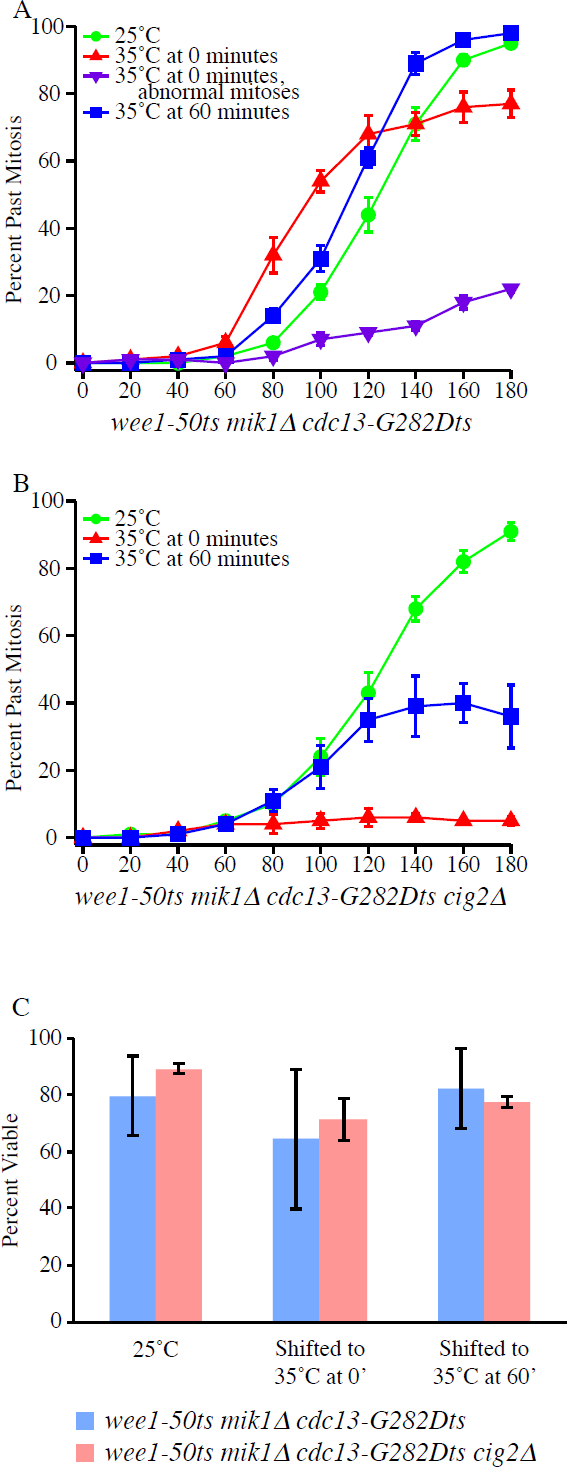
Cig2 can drive viable mitoses. A,B) yFS1004 (*wee1-50ts mik1Δ cdc13-G282D*) (A) and yFS1010 (*wee1-50ts mik1Δ cig2Δ cdc13-G282Dts*) (B) were grown to mid-log phase at 25°C, synchronized in late S phase/early G2 by elutriation, grown at the indicated temperatures and examined every 20 minutes for septation. For *wee1-50ts mik1Δ cdc13-G282Dts* cells, abnormal (multiple or asymmetric) septations were also recorded. Error bars represent mean and standard deviation. n = 3 C) Viability of cells after division. Divided cells from the indicated cultures from A and B were micromanipulated to isolated spots on a YES plate, incubated for three days at 25°C and scored for viability by their ability to form colonies. Error bars represent mean and standard deviation. n = 3.

In contrast to the Cig2-dependence of mitosis early in the cell cycle, cells of both genotypes divide when shifted at 60 minutes after elutriation, in mid G2 after Cig2 levels have declined and Cdc13 levels have increased (Creanor and Mitchison, 1996; Yamano et al., 2000) (Figure 5A,B). This result suggests that, during the temperature shift, Wee1 is inactivated more quickly than Cdc13, allowing Cdc2-Cdc13 complexes to be dephosphorylated and activated to trigger mitosis before Cdc13 loses activity. The fact that only half of *wee1-50ts mik1Δ cig2Δ cdc13-G282Dts* cells divide when shifted to 35°C in mid G2 suggests that in some cells Wee1 activity is lost before Cdc13 activity and cells enter mitosis, while in other cells Cdc13 activity is lost first, causing cells to arrest. The higher percentage of *wee1-50ts mik1Δ cdc13-G282Dts* cells that divide when shifted to 35°C in mid G2 suggests that some Cig2 activity lingers at that point and is able to contribute to mitotic entry, consistent with the results presented in Figure 4.

To directly test if Cig2 is sufficient to provide all essential mitotic functions and to replace Cdc13 in cycling cells, we tested the ability of constitutively expressed Cig2 to recuse the lethality of the *cdc13-G282Dts* allele. We expressed Cig2 from the *nmt81* promoter, which is active in the absence of thiamine (Basi et al., 1993), in a *cdc13-G282Dts* background. *nmt81:cig2 cdc13-G282Dts* cells grow at 36°C, the restrictive temperature for *cdc13-G282Dts* (Figure 6A,B). This rescue of the *cdc13-G282Dts* G2 arrest phenotype can also be seen acutely, in that *cdc13-G282Dts* arrest as unseptated cells when shifted to the restrictive temperature of 36°C, but *cdc13-G282Dts nmt81:cig2* cell, which expressing Cig2 in EMM2 media, continue to divide (Figure 6C). These results demonstrate that Cig2 can provide all of the functions necessary for a viable mitosis in the absence, or near absence, of Cdc13/CDK activity.

**Figure 6.**
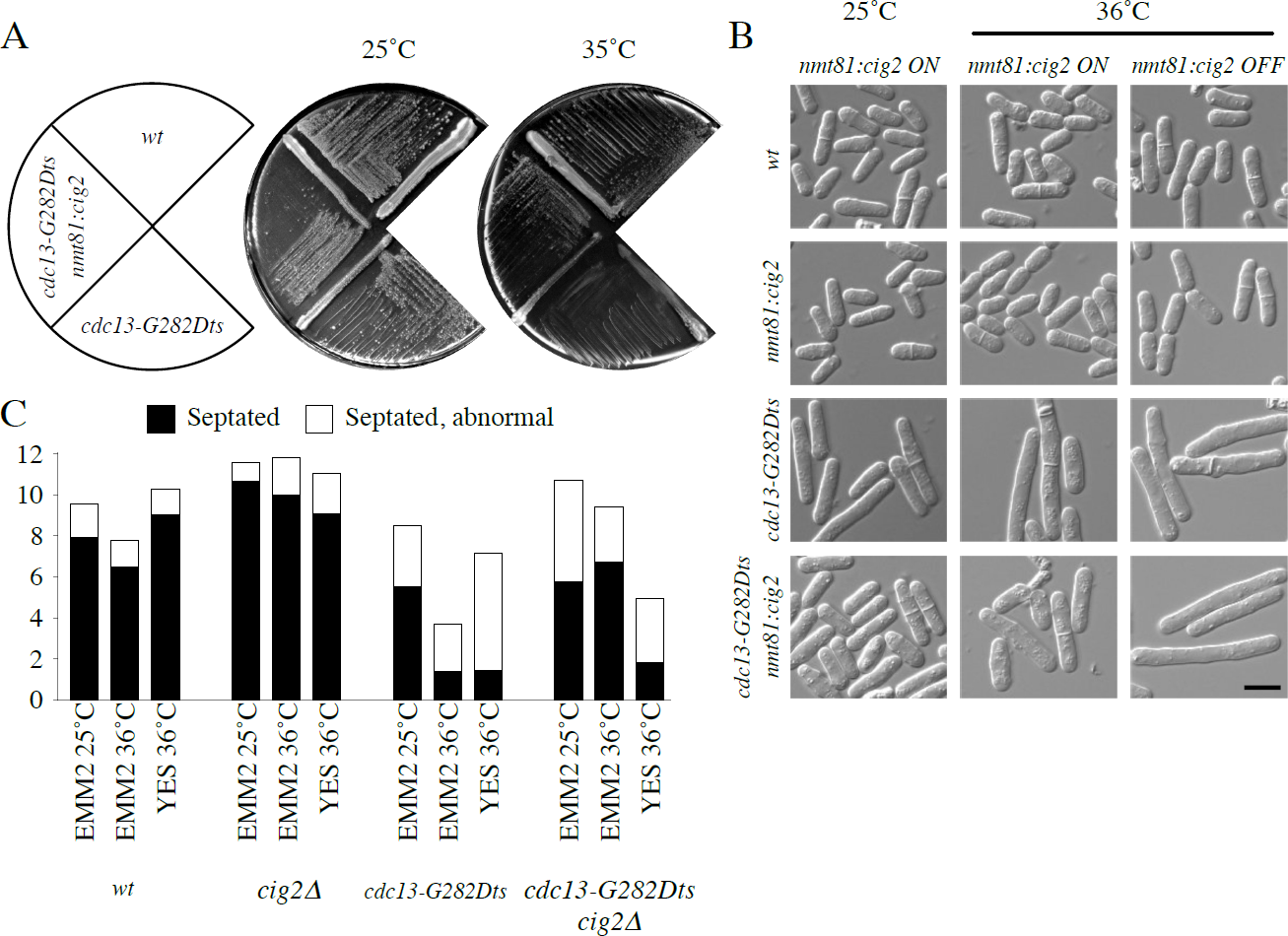
Constitutive expression of Cig2 suppresses loss of Cdc13. A) Plate assay of cell viability. yFS104 (wild type), yFS1032 (*cdc13-G282Dts*) and yFS1048 (*cdc13-G282Dts nmt81:cig2*) cells were struck out on EMM2 plates and grown at the indicated temperatures. B) DIC micrographs of the same strains grown to mid-log phase in liquid EMM2 at 25˚C and incubated at 36°C in EMM2 or YES for 4 hours. The scale bar is 10 *μ*m. C) Cells from the cultures in B were photographed and counted for septation. Abnormal septation includes multiple septa, misshapen septa and asymmetric septation. n > 200.

## Discussion

Cdc13 is the major B-type cyclin in fission yeast, required for entry into mitosis and able to drive S phase in the absence of the other fission yeast B-type cyclins (Booher et al., 1989; Moreno et al., 1989; Fisher and Nurse, 1996). The ability of Cdc13 to regulate an essentially normal cell cycle as the sole cyclin in the cell inspired the quantitative model of cell-cycle regulation, in which varying levels of B/CDK activity trigger various cell-cycle transitions in the proper order (Stern and Nurse, 1996). Although specific expression patterns and substrate-targeting sequences have evolved to optimize specific B-type cyclins to specific cell-cycle transitions, the quantitative model appears to provide the basic mechanism needed to organize a eukaryotic cell cycle (Uhlmann et al., 2011; Fisher et al., 2012). A corollary of the quantitative model is that B-type cyclins that are normally restricted to expression during S-phase (the so-called S-phase cyclins) should be able to drive mitosis if expressed at sufficient levels (Palou et al., 2015).

The ability of Cig2, a fission yeast S-phase cyclin, to drive mitosis is suggested by the fact that it is required for catastrophic mitosis in *wee1-50ts mik1Δ* cells shifted to the restrictive temperature of 30°C (Bueno and Russell, 1993 and Figure 1), at which less than 10% of Wee1 activity remains (Rhind and Russell, 2001). Wee1 and Mik1 are the two Cdc2-Y15 kinases. In their absence, Cdc2 is rapidly dephosphorylated and activated. In G2 cells, this dephosphorylation drives cells into a normal mitosis. However, in cells that have not finished S phase, it drives them into a catastrophic mitosis, in which unreplicated chromosomes are improperly segregated, leading to a cut phenotype and cell death (Lundgren et al., 1991). At 30°C, such catastrophic mitosis requires Cig2, as demonstrated by the viability of *wee1-50ts mik1Δ cig2Δ* cells (Bueno and Russell, 1993 and Figures 1 and 5). Presumably, in the absence of Cig2, the slower accumulation of Cdc13 allows sufficient time between the initiation of S phase and the initiation of mitosis for DNA replication to complete before chromosomes are segregated (Figure 7). Consistent with this conclusion, synchronized *wee1-50ts mik1Δ cig2Δ* cells divide about 1 *μ*m longer than *wee1-50ts mik1Δ* cells (8.8±0.8 *μ*m v. 9.9±0.9 *μ*m, Figure 2)

**Figure 7.**
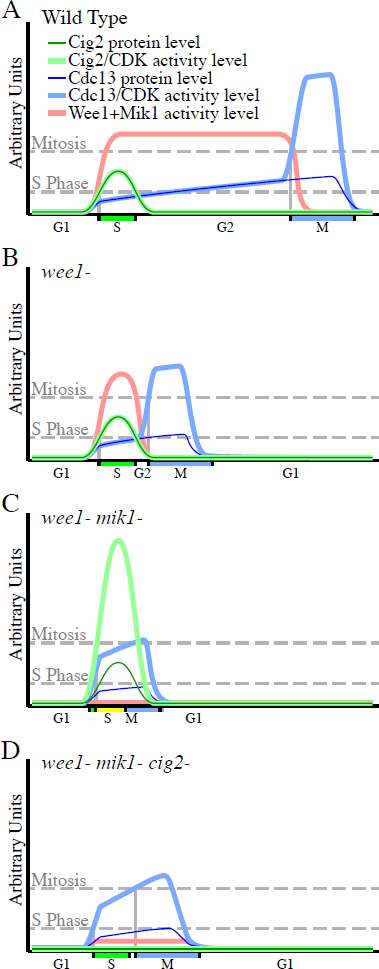
A model for the quantitative regulation of the fission yeast cell cycle. A) In wild-type cells, Cig2 protein is expressed in a brief, symmetric pulse early in the cell cycle (Yamano et al., 2000). Cdc13 gradually accumulates, increasing about two fold during G2 (Creanor and Mitchison, 1996). Mik1 is expressed in a pattern similar to Cig2 (Christensen et al., 2000). Wee1 is expressed at a constant level during G2 (Aligue et al., 1997; Keifenheim et al., 2017), but inactivated by negative feedback from B/CDK activity during mitosis (Tang et al., 1993; McGowan and Russell, 1995). Y15-phosphorylated Cig2 drives S-phase early in the cell cycle (Mondesert et al., 1996). Interacting feedback loops between Cdc13/Cdc2, Wee1 and Cdc25 inactivate Wee1 and Y15-deposhphorylate and activate Cdc2 at the G2/M transition, increases Cdc13/Cdc2 activity about 4 fold (Millar et al., 1991; Rhind and Russell, 1998) and driving cell into mitosis (Rhind and Russell, 2012). B) In the absence of Wee1, Mik1 maintains the Y15 phosphorylation of Cig2/Cdc2 during S phase. Upon the degradation of Cig2 and Mik1, Y15-dephosphorylated Cdc13/Cdc2 is activated and drives cells into mitosis. C) In the absence of Wee1 and Mik1, Cig2 is activated to a level that can drive overlapping S phase and mitosis, leading to mitotic catastrophe. D) In the absence of Wee1, Mik1 and Cig2 (under circumstances in which Cdc13/Cdc2 is not completely active, such as in *wee1-50ts mik1Δ cig2Δ* cells at 30°C, in which ˜10% of Wee1 activity remains (shown), or *wee1-50ts mik1Δ cdc13-RLuc* cells (not shown)), the gradual increase in Cdc13 levels drives S phase before it drives mitosis, leading to a viable cell cycle. Under circumstances in which Cdc13/Cdc2 is completely active, such as in *wee1-50ts mik1Δ cig2Δ* cells at 35°C, Cdc13 is activated quickly enough that S phase and mitosis overlap, leading to mitotic catastrophe (not shown).

Cig2 and Cdc13 appear to cooperate in triggering mitosis. Reduction of either Cig2 or Cdc13 activity can suppress mitotic catastrophe in *wee1-ts mik1Δ* cells at 30°C (Figure 3), and loss of Cig2 enhances the phenotype of the *cdc13-G282D* temperature-sensitive allele (Figure 4). This cooperativity between Cig2 and Cdc13 in promoting mitotic catastrophe allows for the possibility that Cig2 does not drive mitosis itself, but instead simply triggers positive feedback loops between CDK, Cdc25 and Wee1 that hyperactivates low levels of Cdc13, which then goes on to drive mitosis (Zarzov et al., 2002). However, this possibility does not appear to be the case, because Cig2 is sufficient to drive mitosis when Cdc13 is inactivated by *cdc13-G282Dts*, a strong temperature-sensitive allele (Figure 5). In these experiments, when Cdc13 is inactivated during late S/early G2, most cells seem to go through a morphologically normal and viable mitosis (Figure 5). The 20% of cells that go though morphologically abnormal mitoses may do so because they enter mitosis too early, before S phase is complete, leading to a cut phenotype, or because they enter mitosis too late, by which point Cig2 has declined to a level at which it is unable to drive a normal mitosis.

The fact that Cig2 is not sufficient for viability in wild-type cells is presumably due to its expression pattern. Cig2 is expressed in a brief pulse coinciding with S phase (Yamano et al., 2000). It also coincides with the expression of Mik1 (Christensen et al., 2000). Cig2 and Mik1 are concomitantly degraded by SCF ubiquitin-ligase-dependent proteolysis, ensuring that Cig2/CDK is always tyrosine phosphorylated and thus can drive S phase, but can never drive mitosis. However, this model raises the question of why *wee1-50ts cig2Δ* cells are alive. Why does Cdc13/CDK not drive S phase and mitosis concomitantly when it is eventually expressed, leading to a lethal cut phenotype? One possibility is that Cdc13 expression increases slowly enough that the time between triggering S phase and mitosis is sufficient to fully replicate the genome. We invoke this possibility to explain the viability of *wee1-mik1-cig2-*cells (Figure 7D). However, tyrosine phosphorylation is sufficient to delay Cdc13-dependent S phase in cells lacking Cig2 (Mondesert et al., 1996), suggesting that when Mik1 expression decreases in *wee1-50ts cig2Δ* cells at restrictive temperature, Cdc13 activity should quickly rise, triggering both S phase and mitosis in rapid succession. The fact that it does not, and that *wee1-50ts cig2Δ* cells are viable at restrictive temperature (Bueno and Russell, 1993), suggests that Mik1 expression or activity may be regulated by ongoing DNA replication and delay mitosis in *wee1-50ts cig2Δ* cells until S phase is complete (Christensen et al., 2000). Consistent with this hypothesis, *wee1-50ts cig2Δ* cells at restrictive temperature are about 2 *μ*m longer than *wee1-50ts* control cells (Bueno and Russell, 1993).

The fact that Cig2 is able to drive mitosis in the absence of Cdc13 when relieved of inhibitory tyrosine phosphorylation leads us to propose a revised quantitative model of the fission yeast cell cycle (Figure 7). Our model illustrates how Cig2 drives catastrophic mitosis in the absence of Wee1 and Mik1 and why Cdc13 is unable to do so. Although this model lacks the formal rigor of more sophisticated computational models of the fission yeast cell cycle (Novak and Tyson, 1995; Gerard et al., 2015; Keifenheim et al., 2017), it captures cooperation between the cyclins that the mathematically rigorous models currently lack and makes novel, testable predictions, such as the role of Mik1 regulation in delaying mitosis in *wee1-50ts cig2Δ* cells, as discussed above and observed by Coudreuse and Nurse Coudreuse and Nurse (2010 Fig. S19;). The integration of concepts from this model into rigorous computational models will improve our understanding of the details of the fission yeast cell cycle and eukaryotic cell cycles in general.

The ability of Cig2 to drive mitosis in fission yeast also explains the surprising result that cells expressing the Cdc13-Cdc2 single-chain CDK do not require Cdc2 Y15 phosphorylation for viability or size control (Coudreuse and Nurse, 2010). Dephosphorylation of Cdc2 Y15 is the rate-limiting event for entry into mitosis in wild-type fission yeast and otherwise wild-type cells lacking Y15 phosphorylation divide at inviably small sizes (Gould and Nurse, 1989; Lundgren et al., 1991). However, when a strain expressing a translational fusion of Cdc13 and Cdc2 was found to be viable and have proper size control in the absence of Cdc2 Y15, it was proposed that there must be another, Y15-independent, mechanism that restrains Cdc13/CDK activity until cells are large enough for viable mitosis (Coudreuse and Nurse, 2010). Our results suggest that this mechanism is regulation of Cdc13 expression. Specifically, we suggest that the Cdc13-Cdc2 single-chain CDK is not expressed early enough to cause mitotic catastrophe and that, due to the absence of Cig2/CDK activity, mitosis is delayed until sufficient Cdc13-Cdc2 is expressed to drive mitosis (Figure 7). Since the Cdc13-Cdc2 single-chain CDK is expressed from the *cdc13* promoter, it presumably retains both the transcriptional and translational regulation of Cdc13. Although tyrosine phosphorylation is a general mechanism for size control in fungal and metazoan somatic cells (Nurse, 1990; Harris, 2006), cyclin expression appears to regulate cell-division timing in many metazoan embryonic cells and in other eukaryotic domains, such as plants and ciliates (Minshull et al., 1989; Harashima et al., 2013). Therefore, Cdc13-Cdc2 cells may reveal a basal eukaryotic size control mechanism based on size-dependent cyclin expression that has been superseded by CDK tyrosine phosphorylation in fungi and metazoans.

The results presented here support a quantitative conception of cell cycle control. However, qualitative differences in B-type cyclin substrate recognition can clearly play an important role. For instance, although the mitotic B-type cyclin Clb2 (Hu and Aparicio, 2005), or the over expression of its paralog Clb1 (Schwob et al., 1994), is sufficient for S phase and mitosis in budding yeast, over expression of the S-phase B-type cyclin Clb5 is not (Haase and Reed, 1999). This specialization of Clb5 is due in part to a substrate-specific docking motif, which allows it to preferentially phosphorylate S-phase substrates even though it has a relatively low specific activity as a general kinase (Kõivomägi et al., 2011). We therefore favor a model in which the quantitative mechanism of cell cycle control is the ancestral state and remains the fundamental mechanism of eukaryotic cell cycle control. Nonetheless, it has been refined, elaborated and optimized by the addition of qualitative, substrate-specificity layers of regulation, which allow for greater flexibility and cell-type specificity (Uhlmann et al., 2011).

## Material and Methods

The strains used in this study are listed in Table 2. Strains were made and maintained using standard methods (Forsburg and Rhind, 2006). Unless otherwise stated, cells were grown at 25°C in YES (yeast extract plus supplements); *nmt81:cig2* strains were grown it EMM2 (Edinburgh minimal medium, version 2) to induce expression from the *nmt81* promoter. For dilution assays, cells were grown to mid log phase, concentrated to 2x107 cells/ml, 10-or 4-fold serial diluted in a microtiter plate, pinned to YES plates, grown for three days at the indicated temperature, and photographed. For doubling-time assays, cells were grown to mid log phase, diluted to OD 0.1, grown for 48 hours in YES at the indicated temperature, measured for optical density, and diluted ten-fold as necessary to prevent the cultures from exceeding OD 1.0. From these same cultures, cells were photographed using DIC optics on a Zeiss Axioskop 2. Cell lengths were measured in ImageJ (Schneider et al., 2012).

**Table 2.**
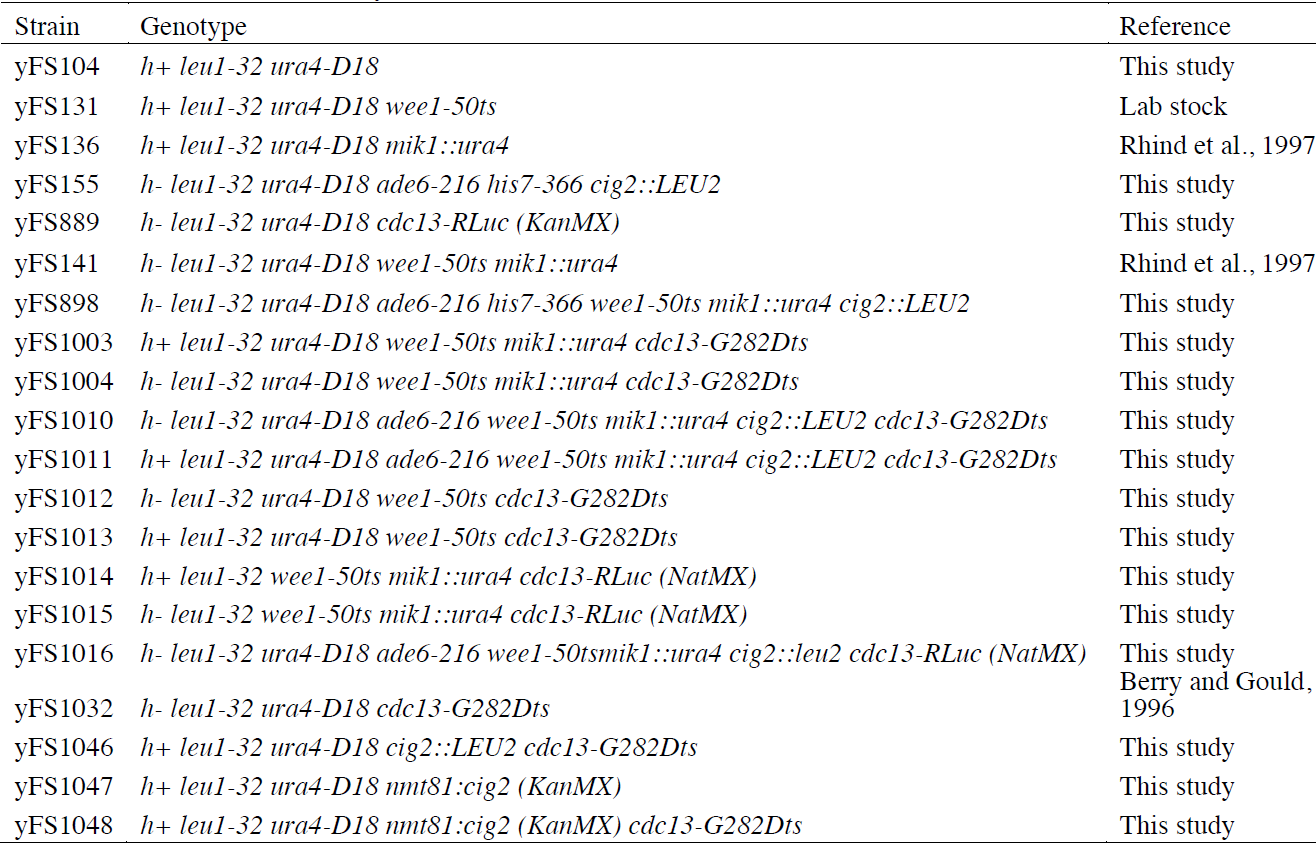
Strains used in this study

The *cdc13-RLuc* allele was made by amplifying pFS346 with DK64 and DK65 and transforming yFS104. The *nmt81:cig2* allele was made by amplifying pFA6a-kanMX6-P81nmt1 (Bahler et al., 1998) with MM71 and MM72 and transforming yFS104. Oligo and plasmid sequences are available by request.

Cells were synchronized by centrifugal elutriation, grown at the indicated temperature and monitored every 20 minutes for septation by bright field microscopy (Willis and Rhind, 2011). Percentage past mitosis was calculated as (# septated cells + # divided cells/2)/(# unseptated cells + # septated cells + # divided cells/2). To determine viability, 64 divided cells were micromanipulated to individual locations on a YES plate, grown three days and scored as viable if a colony was visible.

Time-lapse movies were taken on a DeltaVision OMX with DIC optics using the Freestyle Fluidics approach (Walsh et al., 2017). Briefly, 50 *μ*l of mid log cells were spotted on a lectin-coated (Sigma L1395) glass-bottomed dishes (MatTEK, P35G-1.5-14-C) under 1 cm of FC40 (an oxygen permeable fluorocarbon oil, gift of Peter Cook). Approximately 48 *μ*l of media was removed, leaving a ˜7 mm by 10 *μ*m disk of YES, in which the cells were filmed. Cell lengths were measured in ImageJ (Schneider et al., 2012).

## Acknowledgements

We are grateful to Peter Cook for sharing his Freestyle Fluidics technology with us and to Peter Pryciak and members of the Rhind lab for helpful comments on the manuscript.

